# Sexual dimorphism in the homeostasis of the gut microbiome-immune axis

**DOI:** 10.64898/2026.04.14.718590

**Authors:** Luis Diambra

## Abstract

The gut microbiome is a key regulator of host immunity; however, microbial factors defining basal immune homeostasis remain poorly understood. While large-scale multi-omics have mapped microbial influences on stimulated immune responses, the fundamental dialogue maintaining resting inflammatory tone is often obscured by systemic confounders. In this study, we demonstrate that failing to account for sex leads to a Simpson’s paradox. Using a multivariate framework, we reveal a profound sexual dimorphism in the gut-immune homeostatic axis. In women not using oral contraceptives, we identified consistent homeostatic anchors, including *Bacteroides ovatus, Faecalitalea cylindroides*, and *Firmicutes bacterium CAG:83*, which correlate negatively with TNF-*α* and IL-1*β* levels. This regulatory network is virtually abolished in women using oral contraceptives, revealing an associative silencing where exogenous synthetic hormones uncouple the microbiome from host immune sentinels. Furthermore, taxa such as *Butyricimonas virosa* exhibit sign reversals in cytokine associations depending on hormonal context, highlighting the plasticity of microbial immunomodulation. Our findings establish that basal immune homeostasis is a sex-specific construct and underscore the need for endocrine stratification to accurately decipher the regulatory architecture.

## Introduction

The human gut microbiome represents a dynamic and diverse microbial ecosystem inhabiting the intestinal tract of the human holobiont. Its composition is shaped by a complex interplay between host factor and environmental determinants, most notably dietary habits^1^. Recently, extensive research has elucidated the critical role of the gut-organ axes, highlighting the intricate links between the microbiota and various immune^2^, metabolic^3^, and neurological traits^4,5^ in both health and disease. Under physiological conditions, gut-derived metabolites play a fundamental role in maintaining host homeostasis, facilitating nutrient processing, supporting crosstalk across the gut-brain axis^6^, and orchestrating immune signaling^7^.

The human body is not in a state of zero inflammation but rather in a basal state of controlled inflammation, largely orchestrated by signals from the gut^8^. The human gut microbiome is essential for immune defense, primarily through the precise modulation of cytokine networks^9^. The basal modulation of inflammation constitutes a complex, bidirectional communication axis where both components regulate each other in a dynamic dialogue^10^; any disruption to this interaction can precipitate dysbiosis and systemic inflammation. The basal modulation of inflammation by the gut microbiota in healthy subjects is one of the current topics that attracts the attention of many researchers in the field^11–14^.

The gut is in constant communication with the systemic immune system through microbial metabolites and the regulation of intestinal permeability. On one hand, the microbiome acts as a central rheostat of the immune system, dictating the production and release of specific signaling molecules. For instance, certain commensal bacteria produce muropeptides and short-chain fatty acids (SCFAs, such as butyrate, propionate, and acetate) that induce anti-inflammatory cytokines, thereby dampening immune activation and inhibiting the synthesis of pro-inflammatory mediators^15,16^. Conversely, bacterial cell wall components are recognized by pattern recognition receptors, triggering the release of TNF-*α*, IFN-*γ*, IL-6, and IL-1*β* to counter potential pathogens^17^. Thus, the host’s cytokine profile exerts selective pressure on the composition and behavior of microbial communities. Through these mechanisms, cytokines preserve ecological stability by preventing opportunistic species from dominating the niche in a basal state of controlled inflammation^17^. However, if the intestinal barrier is weakened, small amounts of bacterial products can enter the systemic circulation, raising the baseline inflammatory tone. A persistent excess of pro-inflammatory cytokines can perturb the intestinal environment, leading to a loss of microbial diversity and the expansion of pathobionts, a condition known as dysbiosis^18^. Thus, structural disruption of the gut microbiota can drive the progression of several chronic diseases, including inflammatory bowel disease, metabolic syndrome, and neurodegenerative diseases, and can disrupt local and global immune responses.

Given the critical role of the microbiota in modulating immune responses and the conexion with health and well-being, this axis has become a focal point of contemporary research. A important milestone in this field is the Human Functional Genomics Project (HFGP), a comprehensive initiative designed to unravel the mechanisms underlying inter-individual variation in immune responses^19–21^. Specifically, the HFGP provides an unprecedented framework to investigate the intricate relationships between gut microbial composition and systemic inflammatory cytokine levels on immune homeostasis of healthy individuals^21^. By integrating multi-omic data from a well-characterized cohort of almost 500 healthy volunteers of Western European ancestry, this project enables the identification of key factors –inlcuding host genetics (SNPs) intrinsic variables (age, gender, and body mass index (BMI)) and environmental determinants as wheather and oral contraceptives (OC) use, and the gut microbiome– that collectively drive the inter-individual variability in the production of three monocyte-derived cytokines (IL-1*β*, TNF-*α*, IL-6) and three lymphocyte-derived cytokines (IFN-*γ*, IL-17, IL-22).

Previous reports from this cohort have highlighted a strong influence of age and sex on cytokine production independent of the microbiome^19^. Meanwhile, a sister study identified several microbiome-cytokine interaction patterns; however, these associations were established using univariate Spearman-rank correlations, that is, independent of the intrinsic variables of individuals^21^. This preliminary analysis did not account for the confounding effects of age and sex, despite the evidence that these factors significantly influence cytokine levels. The reliance on univariate analyses not only risks overlooking subtle microbial influences but also leaves the data susceptible to Simpson’s paradox^22^, as we will show later. To address these limitations, we perform a sex-stratified analysis implementing a multivariate residual-based correlation framework. An overview of data and study design used here are sketched in Fig. 1. This refined approach allowed us to uncover a profound sexual dimorphism in basal immune homeostasis and demonstrates how oral contraceptives decouple the gut-immune axis, effectively silencing the microbial signals that otherwise sustain host inflammatory tone.

**Figure 1.**
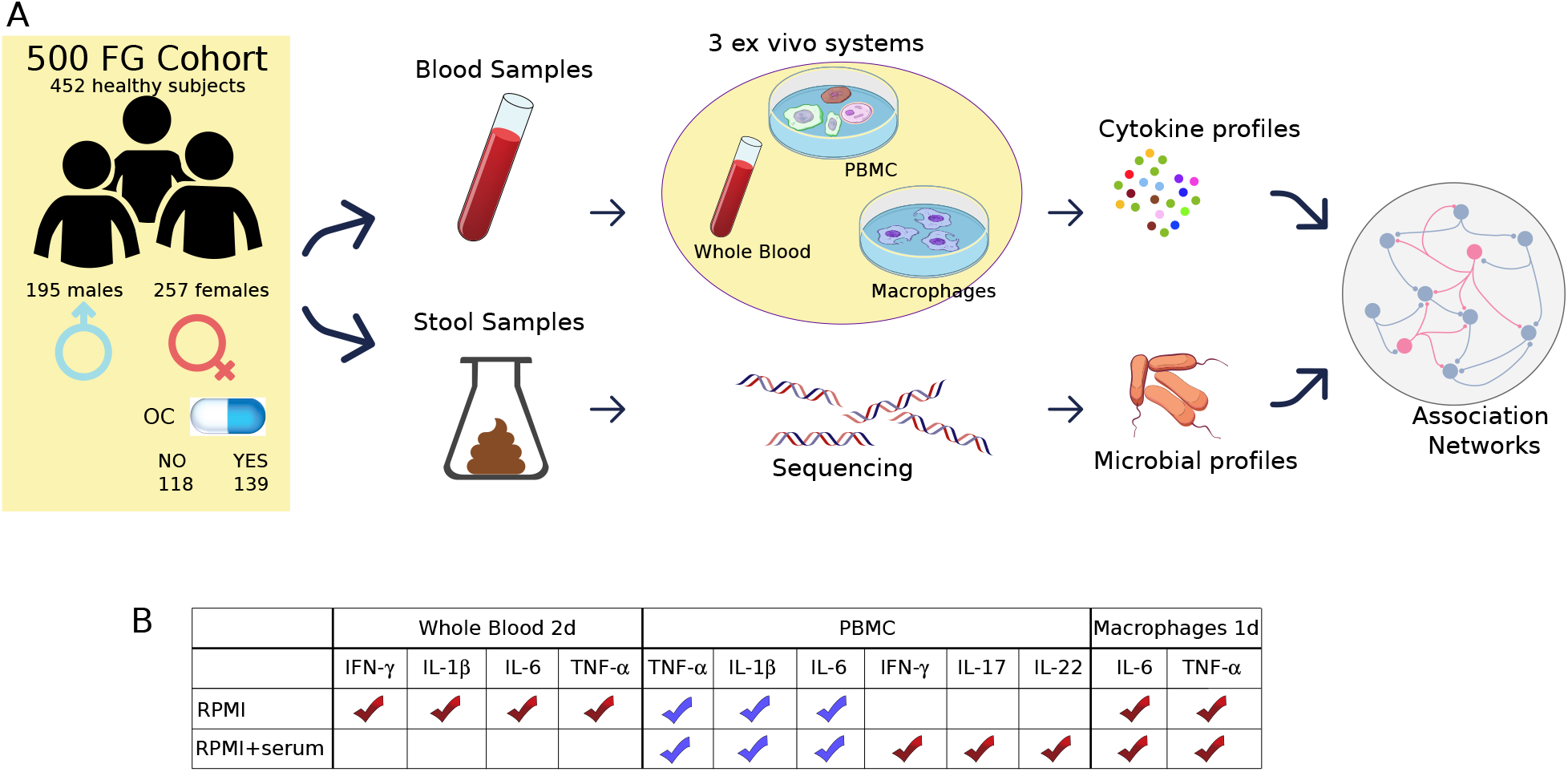
Schematic overview of the data used in this study. (A) Microbial and cytokine profiles available publicly from 500FG cohort were integrated. It comprising 452 healthy subjects after stringent exclusion criteria. The cohort was stratified by sex and, within the female subgroup, by oral contraceptive (OC) use. Cytokines were measured across three different *ex vivo* systems: whole blood, peripheral blood mononuclear cells (PBMCs), and monocyte-derived macrophages, and were associated with microbial profiles to determine the baseline immune associative network at different levels of cellular complexity. (B) Cytokines were assessed under two different basal conditions: RPMI and RPMI+serum. Not all cytokines were assessed in the three systems and in both conditions. The table specifies the cytokines measured according to the condition and systems: in the whole blood system four cytokines were evaluated for RPMI condition; in the PBMC systems a temporal distinction was made between acute (1-day, blue) and long-term (7-days, red) responses; in the macrophage system the focus was on IL-6 and TNF-*α* in both basal conditions.

## Results

### Simpson’s paradox in the microbiome-cytokine interactions context

The univariate analyses, when outcomes depend on several confounding variables, can overestimate some relationships while underestimating others, leaving the data susceptible to Simpson’s paradox. To identify and mitigate spurious associations between the gut microbiome and cytokine levels, we implemented a residual-based modeling framework across two distinct scenarios: (1) a global analysis of the entire cohort, incorporating sex as a covariate alongside age, BMI, and OC use; and (2) a sex-stratified analysis, where age, BMI, and OC use (specifically for the female subgroup) were treated as covariates. This dual-scenario approach was instrumental in detecting instances of Simpson’s paradox, where aggregate trends contradicted the associations found within individual strata. Figure 2 illustrates the Spearman-rank correlation coefficients (*R*) between cytokine residuals and microbial abundances under basal conditions (RPMI and RPMI+serum) for two *ex vivo* systems: whole blood (Fig. 2A) and peripheral blood mononuclear cells (PBMCs, Fig. 2B). Associations were corrected using the Benjamini-Hochberg method, where grey boxes indicate significant relationships in the aggregate analysis (FDR*<* 0.01), overlaid with the corresponding *R* values from the sex-stratified analysis (blue disks for males and red disks for females). While certain microbe-cytokine associations persisted after sex stratification, such as *Butyricimonas virosa* with IL-1*β* in whole blood and *Lawsonibacter asaccharolyticus* with IL-1*β* in PBMCs, many other links were identified as spurious. For instance, although *A. finegoldii, C. leptum*, and *B. uniformis* exhibited significant positive correlations with IL-1*β* in the aggregate whole blood analysis, these associations disappeared upon stratification (Fig. 2A). Furthermore, clear cases of sign reversal were observed, most notably between *A. finegoldii* and TNF-*α* (Fig. 2A), with additional instances showing similar trends in macrophages *ex vivo* system (see Fig. S1) and at lower significance levels (see Table S1 for FDR *<* 0.05). Furthermore, we identified several instances of sex-dependent divergence in microbe-cytokine associations, where opposing trends were observed between males and females. Notable examples include the association between *O. sp. CAG:241* and TNF-*α* (Fig. 2A and B), and *A. muciniphila* with IL-6. By adopting this dual-scenario approach, we aimed to decouple confounding effects and unmask the intrinsic biological signals. Our results demonstrate that the omission of sex-specific adjustments in microbiome-cytokine studies can lead to misleading qualitative conclusions driven by Simpson’s Paradox. Consequently, in the following sections, we prioritize the stratified, covariate-adjusted analysis, as it provides the most biologically accurate representation of these interactions within a sexual dimorphism framework.

**Figure 2.**
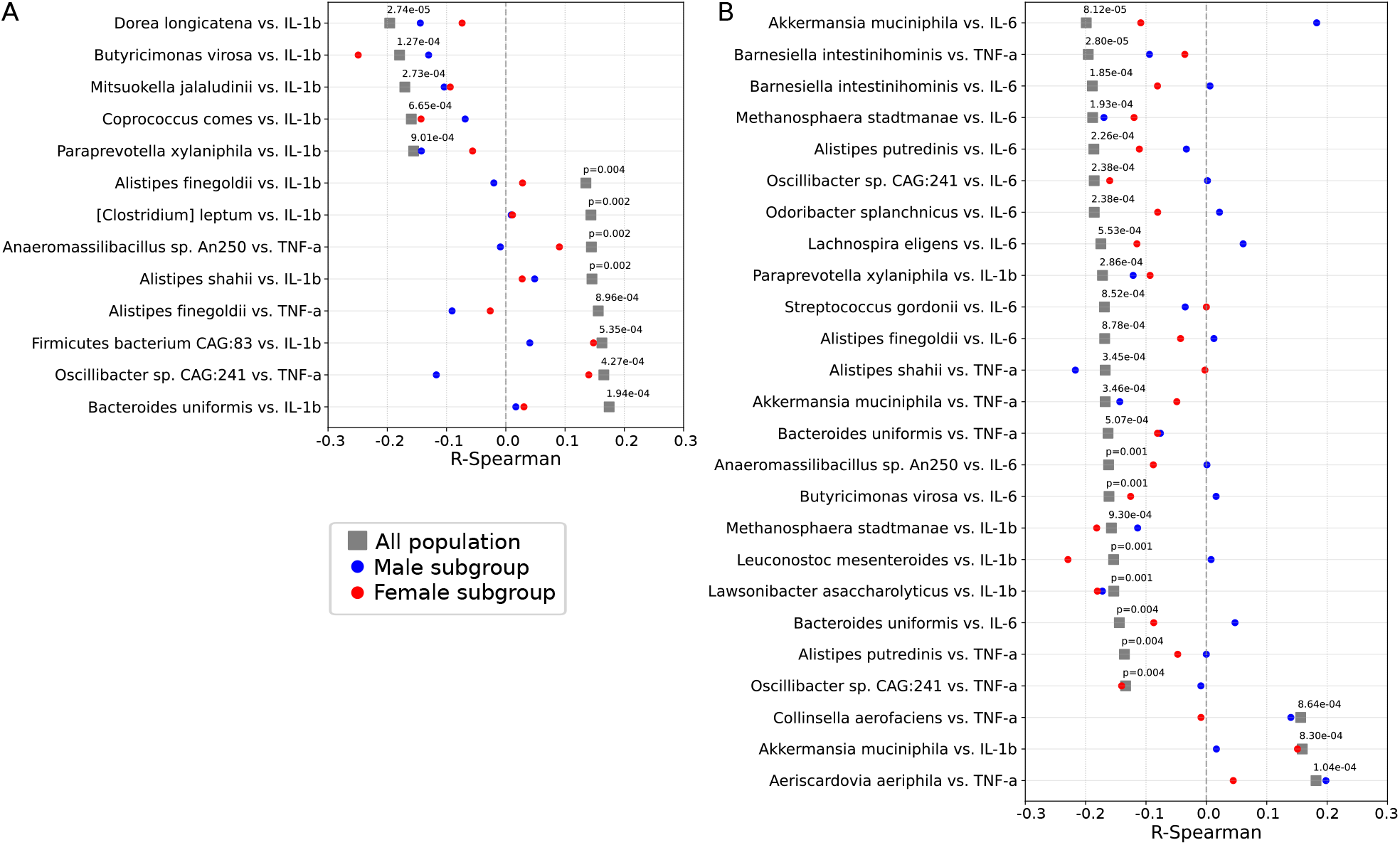
Significant associations between microbial species and cytokine residuals on global analysis. Forest plot of Spearman-rank correlation coefficients are shown for non-stimulated controls (RPMI and RPMI+serum indicated by *) across two *ex vivo* systems: (A) whole blood and (B) PBMCs. Residuals were derived from OLS models adjusting for sex, age, BMI, and oral contraceptive use. Significance was determined using Benjamini-Hochberg FDR correction, where grey boxes indicate significant relationships in the aggregate analysis (FDR *<* 0.01), overlaid with the corresponding *R* values from the sex-stratified analysis (blue disks for males and red disks for females). For similar forest plot for monocyte-derived macrophages see Fig.S1. Only species detected in > 3% of all samples were included in the analysis.

### Links between the gut microbiome and cytokine levels characterizing host immune homeostasis

By adopting the stratified approach, we aim to uncover a more realistic representation of the microbiome-cytokine interactions underlying the host immune homeostasis. In this sense we compute Spearman-rank correlation coefficients (*R*) between cytokine residuals and microbial abundances. The results obtained for three *ex vivo* cellular systems (whole blood, PBMC, and monocyte-derived macrophages) under two basal conditions (RPMI and RPMI+serum) are summarized in Fig. 3 using an undirected association network, and commented on in the subsections below. To complement these networks, we also provided forest plots (Supplementary Figures S2-S4) that detail the correlation coefficients (*R*) for each significant association and tables (see Table S2 at significance levels FDR *<* 0.05). The forest plots display not only the significant *R* value for one sex but also the corresponding coefficient for the opposite sex, regardless of its significance. This side-by-side comparison explicitly highlights the sexual dimorphism and cases of sign reversal, providing a more granular view of the divergent microbial-host interactions.

**Figure 3.**
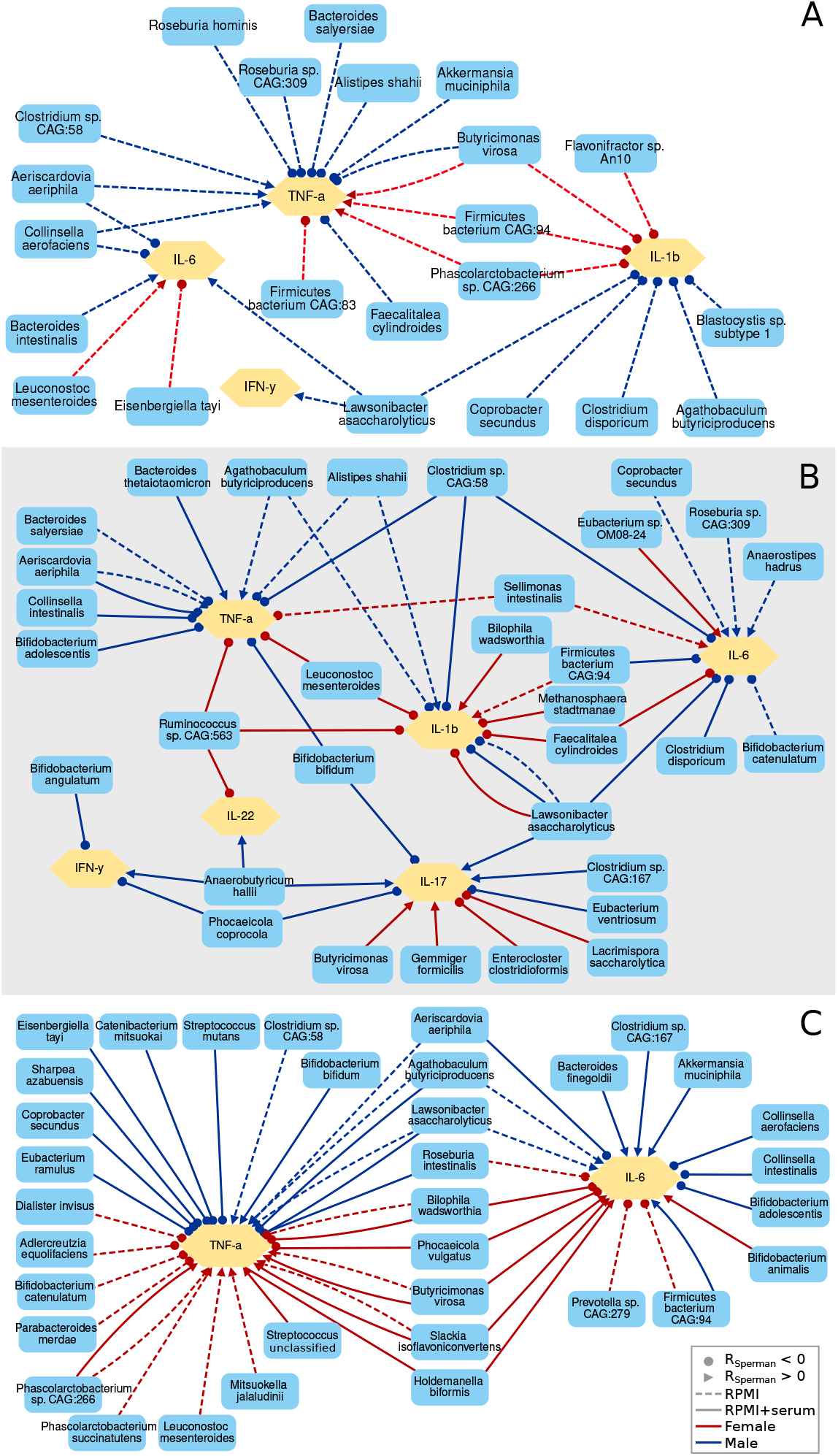
Sex-stratified analysis. Undirected association network depict significant relationships between microbial species and cytokine residuals obtained the under basal conditions (RPMI and RPMI+serum) for three *ex vivo* cellular systems: whole blood (A), PBMCs (B) and monocyte-derived macrophages (C). Microbial species (blue nodes) and cytokines (yellow nodes) are linked by edges representing significant relationships. Edge colors distinguish between the sexes, with blue edges indicating associations in males and red edges indicating those in females. Dashed and solid edges indicate the RPMI and the RPMI+serum conditions, respectively. To denote the nature of the association without implying a regulatory direction, edges are capped with specific markers: disks represent negative correlations (*R <* 0), while arrows represent positive correlations (*R >* 0). Residuals were derived from OLS models adjusting for age, BMI, and OC use for each gender subgroups. Significance was determined using Benjamini-Hochberg FDR correction (*FDR <* 0.05). Only species detected in > 3% of all samples were included in the analysis.

#### Whole blood system

In the whole blood system, four cytokines were assessed under the RPMI condition, identifying 30 significant associations (*FDR <* 0.05). Several taxa emerged as potential drivers of immune homeostasis through consistent anti-inflammatory associations (Fig. 3A). However, these protective signals frequently exhibited marked sexual dimorphism. A notable example of this sexual divergence was observed among species negatively correlated with IL-1*β* ; specifically, no single taxon maintained a significant link with IL-1*β* across both sexes, suggesting that the microbial regulation of this cytokine is highly sex-specific. Several well-recognized homeostatic anchors, including *Alistipes shahii* (*R* = −0.26), *Roseburia hominis* (*R* = −0.21), and *Akkermansia muciniphila* (*R* = −0.20), were negatively associated with TNF-*α* levels in males^23–25^. Similarly, *Faecalitalea cylindroides, Roseburia sp. CAG:309*, and *Bacteroides salyersiae* displayed negative correlations with TNF-*α* in the male cohort. Furthermore, both the protist *Blastocystis sp. subtype 1* and the bacterium *Lawsonibacter asaccharolyticus* showed significant negative correlations with IL-1*β* in males (*R* = −0.21 and *R* = −0.27, respectively). Interestingly, *L. asaccharolyticus* also exhibited a positive modulation of IFN-*γ* and IL-6. While these cytokines are typically associated with acute inflammatory responses, their upregulation under basal conditions by *L. asaccharolyticus* may suggest a role in enhancing immune surveillance and maintaining epithelial homeostasis rather than promoting chronic inflammation^26,27^. The most prominent case of sexual dimorphism and sign reversal was observed for *Butyricimonas virosa*. While it displayed a clear anti-inflammatory profile in males –negatively correlating with TNF-*α* (*R* = −0.22)– it showed a discordant, positive association with pro-inflammatory TNF-*α* (*R* = 0.20) in females. This divergence underscores the necessity of sex-stratified analyses to correctly interpret the immunomodulatory potential of the human gut microbiota. The associated forest plot is reported in Fig.S2.

#### PBMC system

In the PBMC system, the six cytokines were assessed under both RPMI (dashed lines) and RPMI+serum (solid lines) conditions. The removal of granulocytes and plasma factors allowed for a more granular view of the mononuclear response. Fig. 3B depicts 14 significant associations that were identified under the RPMI condition, while 37 significant associations were identified under the RPMI+serum condition (at *FDR <* 0.05). Under RPMI condition, we observed potent anti-inflammatory signatures in males. For instance, *Agathobaculum butyriciproducens* and *L. asaccharolyticus* showed strongest negative correlations with IL-1*β* (*R* = −0.31 and *R* = −0.29, respectively), while *Bifidobacterium catenulatum* were consistently associated with reduced IL-6 levels (*R* = −0.29). The RPMI+serum condition was essential for capturing adaptive immune signals. As noted previously, significant associations with IFN-*γ*, IL-17, and IL-22 only emerged in this supplemented environment, which supports long-term lymphocyte viability. Here, *Anaerobutyricum hallii* and *L. asaccharolyticus* appeared to promote a healthy ‘priming’ of Th17 and Th22 responses.

A remarkable female-specific signature was identified for *Ruminococcus sp. CAG:563*. In the female cohort, this taxon exhibited a targeted inhibitory profile, correlating negatively with two primary pro-inflammatory markers, IL-1*β* and TNF-*α*, as well as with IL-22. While IL-22 is essential for epithelial repair, its downregulation alongside acute inflammatory cytokines suggests that *Ruminococcus sp. CAG:563* promotes a state of immune quiescence in the female gut, potentially reducing unnecessary mucosal activation. In males, similar multiple cytokine suppression in males is exerted by *Clostridium sp. CAG:58*, highlighting the sex-specific nature of this commensal’s role in host homeostasis. Sexual dimorphism was again evident through specific taxa. In females, *Firmicutes bacterium CAG:94* showed a positive correlation with IL-1*β* (*R* = 0.18), contrasting with the male-specific anti-inflammatory patterns. Furthermore, *Leuconostoc mesenteroides* exhibit a significant negative link with IL-1*β* (*R* = −0.24) and TNF-*α* (*R* = −0.17) in female, suggesting that its immunomodulatory effect in women may be more pronounced during prolonged exposures or in the presence of systemic factors. The associated forest plot is reported in Fig.S3.

#### Macrophage system

In the monocyte-derived macrophages system, only two cytokines (TNF-*α* and IL-6) were assessed under both basal conditions. Fig. 3C depicts 20 significant associations that were identified under the RPMI (dashed lines) condition, while 31 significant associations were identified under the RPMI+serum condition (solid lines) at FDR *<* 0.05. The macrophage system provided the highest level of cellular resolution, revealing that sex-specific microbial sensing is intrinsically programmed in the myeloid lineage. In the RPMI+serum condition, we confirmed the male-specific association of *Akkermansia muciniphila* with IL-6 (*R* = 0.27), a link that was absent in females and in serum-free conditions. This underscores the necessity of systemic factors in mediating Akkermansia-host communication. Distinctive patterns also emerged for *L. asaccharolyticus* in males; while it initially showed a positive correlation with IL-6 and TNF-*α* in RPMI, the addition of serum flipped the association with TNF-*α* to negative (*R* = −0.26), suggesting a serum-dependent anti-inflammatory switch. In contrast, the female macrophage landscape was characterized by strong inhibitory links from taxa such as *Bilophila wadsworthia* (*R* = −0.29) and *Bifidobacterium catenulatum* (*R* = −0.19) toward TNF-*α*. Notably, the broad multi-cytokine suppression previously observed for *Ruminococcus sp. CAG:563* in female PBMCs was not replicated in the isolated macrophage model. This indicates that its immunomodulatory effects are not exerted through direct myeloid inhibition but rather through complex cellular cross-talk within the mononuclear cell population. In addition to these findings, our data highlighted the importance of dietary-linked commensals in shaping macrophage polarities. In males, the presence of serum appeared to be a critical switch for the immunomodulatory activity of *Agathobaculum butyriciproducens*; while it showed a positive association with TNF-*α* in RPMI, this link became significantly negative (*R* = −0.22) in the serum-supplemented condition. Furthermore, several female-specific positive links were identified in the presence of serum, most notably for *Holdemanella biformis* and *Slackia isoflavoniconvertens*, both of which were associated with both cytokines. Further, the pro-inflammatory trend of *Butyricimonas virosa* in the female cohort persisted, showing a positive correlation with TNF-*α* in the isolated macrophage model (*R* = 0.16 and *R* = 0.22). This consistency across whole blood and isolated cell systems reinforces the hypothesis that the sex-specific response to certain gut taxa is a robust biological feature, independent of the complexity of the experimental environment. The associated forest plot is reported in Fig.S4.

### Modulating impact of oral contraceptive use on women’s immune homeostasis

Oral contraceptives (OCs) are known to elevate baseline levels of C-Reactive Protein (CRP)^28,29^, a marker of systemic inflammation that may mask the anti-inflammatory links of specific bacterial taxa. Consequently, OC use represents a significant confounding factor in our female cohort, suggesting that the observed microbial-immune associations are nuanced by the host’s exogenous hormonal status. To decouple these effects, we extended our stratified approach to account for OC use. Fig. 4 illustrates this secondary stratification, displaying Spearman-rank correlation coefficients (*R*) between cytokine residues and basal microbial abundance across the whole blood system. Notably, while all associations in the NOC group retain the magnitude and direction consistent with the pre-stratified results, most of these relationships are markedly attenuated in the OC cohort. Furthermore, we identified two instances of opposite polarities involving *Butyricimonas virosa* and *Firmicutes bacterium CAG:94* in their association with IL-1*β* . This divergence suggests that synthetic hormonal intake potentially disrupts or even rewires the endogenous cross-talk between the gut microbiota and the immune system. Despite the limitations in sample size and statistical power introduced by OC-stratification, this approach revealed novel, significant microbial-immune associations that were previously masked by the host’s exogenous hormonal status. In Fig. 5, we highlight these ‘hidden’ links in the three *ex vivo* systems, which only reach significance within the non-OC users (NOC) cohort. Our analysis reveals many more interactions in NOC women’s group compared to the men’s cohort. The emergence of these associations suggests that the physiological hormonal fluctuations of the natural cycle may be necessary to manifest certain immunomodulatory effects of the gut microbiota, which are otherwise suppressed or altered by synthetic steroid intake.

**Figure 4.**
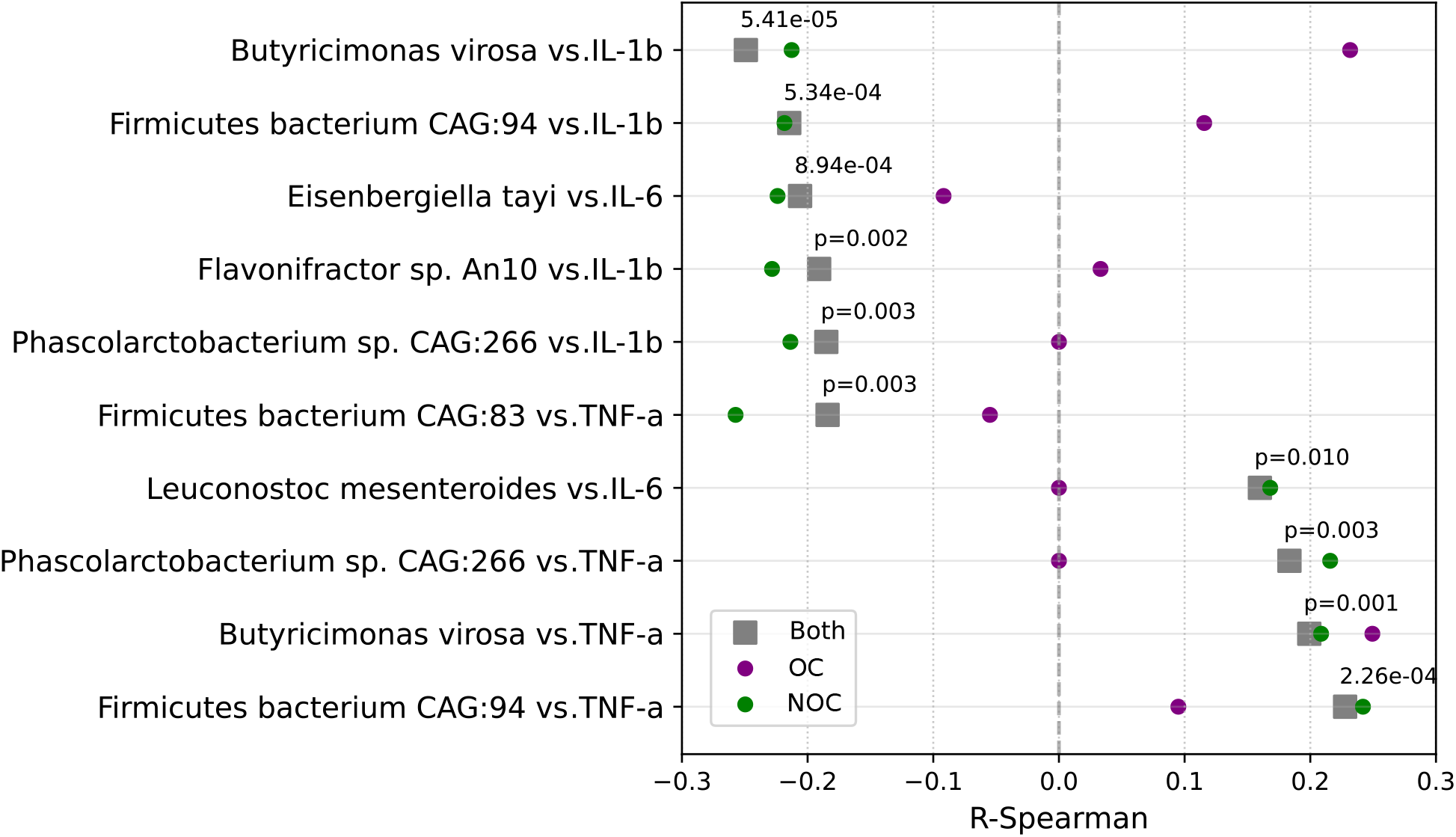
Significant associations between microbial species and cytokine residuals on female cohort. Forest plot of Spearman-rank correlation coefficients are shown for the whole blood *ex vivo* system under RPMI condition. Significance was determined using Benjamini-Hochberg FDR correction, where grey boxes indicate significant relationships in the whole female cohort (FDR *<* 0.05), overlaid with the corresponding *R* values from the OC-stratified analysis: green disks for non-OC users (NOC) cohort and purple disks for OC users (OC). Only species detected in > 3% of all samples were included in the analysis.

**Figure 5.**
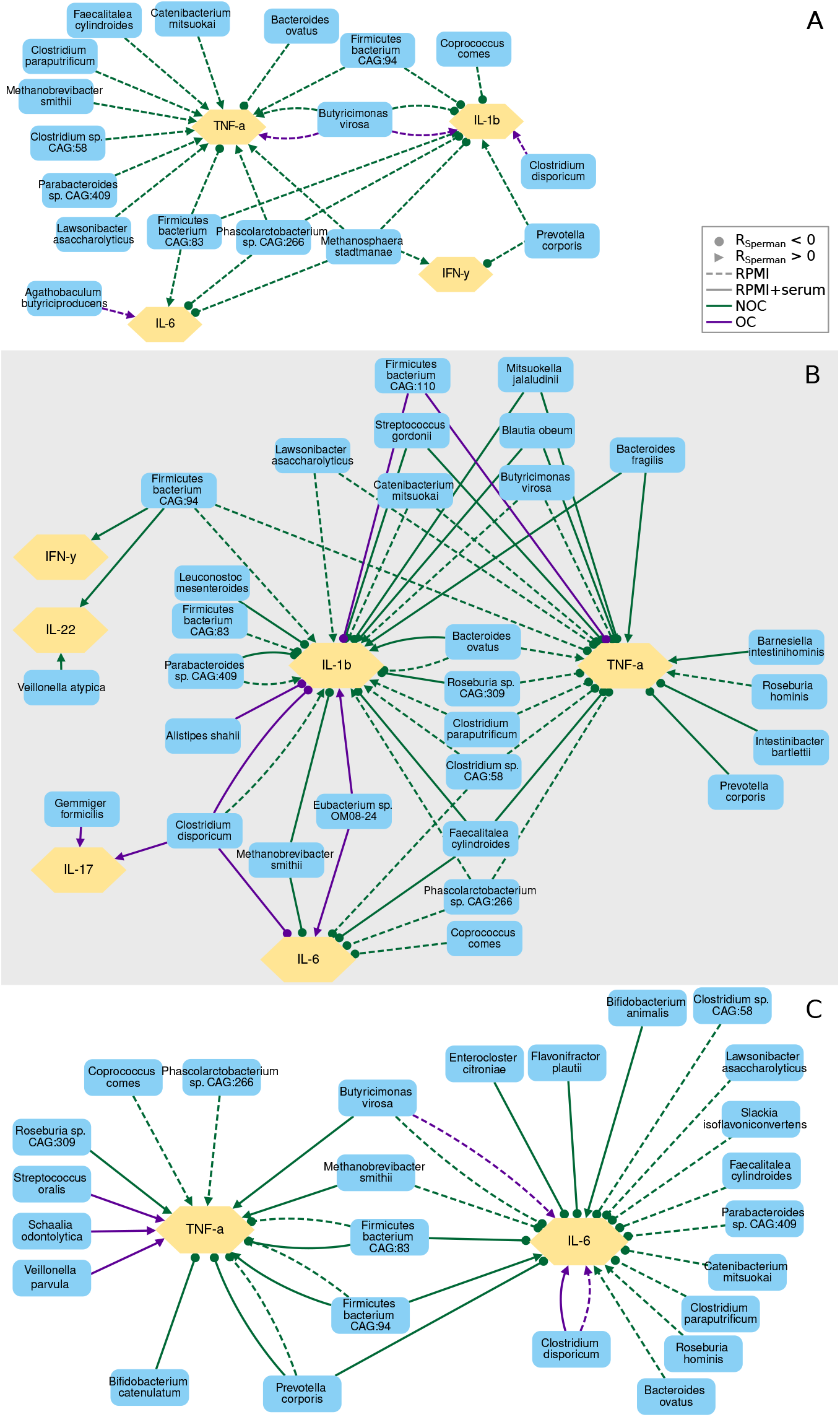
OC-stratified analysis in the women cohort. Undirected association network depict significant relationships between microbial species and cytokine residuals obtained the under basal conditions (RPMI and RPMI+serum) for three *ex vivo* cellular systems: whole blood (A), PBMCs (B) and monocyte-derived macrophages (C). Microbial species (blue nodes) and cytokines (yellow nodes) are linked by edges representing significant relationships. Green edges represent the cohort not using OCs (NOC), while purple discs represent OC users. Dashed and solid edges indicate the RPMI and the RPMI+serum conditions, respectively. To denote the nature of the association without implying a regulatory direction, edges are capped with specific markers: disks represent negative correlations (*R* < 0), while arrows represent positive correlations (*R* > 0). Residuals were derived from OLS models adjusting for age and BMI, for each subgroups. Significance was determined using Benjamini-Hochberg FDR correction (*FDR* < 0.05). Only species detected in > 3% of all samples were included in the analysis. Forest plots reporting these results are shown in Fig.S5, Fig.S6 and Fig.S7.

#### Whole blood system

The stratification of the female cohort by oral contraceptive (OC) use reveals a strong ‘associative silencing’ in the gut-immune axis. While the NOC group displays a rich and complex landscape of 25 significant microbial-cytokine links, the OC group shows a stark reduction to only 4 associations, all of which are exclusively positive. This suggests that exogenous synthetic hormones may override the subtle, metabolite-driven basal modulation of inflammation, effectively masking the contribution of the microbiota to host homeostasis. A pivotal finding in the NOC group is the role of *Bacteroides ovatus*, which emerged as a major homeostatic anchor with the strongest negative correlation with TNF-*α* (*R* = −0.31). This aligns with its known capacity to produce propionate, a SCFA that suppresses pro-inflammatory cascades. However, this association is entirely absent in the aggregate analysis and in women taking OCs, suggesting that synthetic progestins and estrogens might alter the responsiveness of circulating leukocytes to propionate-mediated signaling. The most striking case of sign reversal and functional divergence was observed for *Butyricimonas virosa*. In the NOC cohort, this butyrate-producer exhibited a classic anti-inflammatory signature by negatively correlating with IL-1*β* (*R* = −0.21). In stark contrast, within the OC group, this relationship flipped to a positive correlation (*R* = 0.23), while its association with TNF-*α* also remained positive (*R* = 0.25). This functional reversal suggests that the hormonal environment induced by OCs may not only mask microbial signals but also potentially recalibrate how the immune system interprets metabolites from beneficial taxa, turning a typically tolerogenic signal into a pro-inflammatory one. Furthermore, the NOC group highlighted the influence of the unclassified *Firmicutes bacterium CAG:83*, which appears to act as a central hub for cytokine modulation. It showed a dual profile: while positively correlating with IL-1*β* (*R* = 0.27) and IL-6 (*R* = 0.24), it maintained a significant negative link with TNF-*α* (*R* = −0.25). This nuanced regulation, promoting surveillance cytokines while dampening systemic necrosis factors, is lost in the OC group, further supporting the hypothesis that OCs decouple the microbiota from its role in fine-tuning the basal inflammatory tone. Finally, the inclusion of methanogenic archaea in the NOC network, such as *Methanosphaera stadtmanae* (negatively correlated with IL-1*β* and IL-6), underscores that the homeostatic dialogue extends beyond bacteria. The fact that all archaeal associations vanish under OC use points to a systemic breakdown of the ‘holobiont dialogue’ in the presence of exogenous hormonal interference.

#### PBMC system

The PBMC model provides the most striking evidence of hormonal interference in the gut-immune axis. In NOC subgroup, we observed a highly orchestrated network where taxa such as *Faecalitalea cylindroides* and *Clostridium sp. CAG:58* act as regulatory pivots. For instance, *F. cylindroides* exhibited a consistent inhibitory profile across multiple cytokines, including IL-1*β* (*R* = −0.28), IL-6 (*R* = −0.30), and TNF-*α* (*R* = −0.32). Similarly, *Clostridium sp. CAG:58* emerged as a potent regulator, showing strong inverse associations with TNF-*α* (*R* = −0.33) and IL-1*β* (*R* = 0.32). Interestingly, the PBMC data highlighted the specificity of certain taxa: while *Bacteroides ovatus* is linked with IL-1*β* (*R* = −0.31 in RPMI), it showed a divergent trend with TNF-*α* in this specific cellular environment, suggesting that the microbial modulation of TNF-*α* might be partially dependent on granulocyte-mediated pathways present in whole blood but absent in PBMCs. Remarkably, the quasi-total absence of significant associations in the OC group, with the exception of *Clostridium disporicum* and *Firmicutes bacterium CAG:110*, reinforces the hypothesis that synthetic steroids exert a dominant systemic override on the mononuclear-microbiome dialogue.

#### Macrophages system

The macrophage model, representing the primary orchestrators of tissue inflammation, provided the final evidence of the decoupling effect induced by oral contraceptives. In the NOC cohort, macrophages exhibited a finely-tuned regulatory network where both TNF-*α* and IL-6 were under significant microbial influence. Notably, *Firmicutes bacterium CAG:83* reinforced its role as a consistent homeostatic anchor across all tested systems (whole blood, PBMC, and macrophages), displaying a significant negative correlation with TNF-*α* (*R* = −0.285). This underscores the species’ potential as a universal modulator of the innate immune response in a natural hormonal context. The regulation of IL-6 in NOC macrophages revealed a consortium of anti-inflammatory taxa. Species such as *Catenibacterium mitsuokai* (*R* = −0.27), *Clostridium paraputrificum* (*R* = −0.26), and *Slackia isoflavoniconvertens* (*R* = −0.26) were significantly associated with lower basal IL-6 production. Furthermore, *L. asaccharolyticus* maintained its beneficial profile by negatively correlating with IL-6 (*R* = −0.24), consistent with its previous associations with systemic health. Intriguingly, *Roseburia hominis* showed a positive correlation with IL-6 (*R* = 0.26) in NOC macrophages, suggesting that in this specific cellular compartment, certain butyrate-producers might support a basal level of surveillance signaling without triggering systemic inflammation. However, all these intricate associations –both inhibitory and stimulatory– were completely absent in the OC group. The failure of microbial signals to correlate with macrophage output in OC users suggests that synthetic steroids induce a state of ‘associative silencing’ to commensal metabolites, centralizing immune control away from the microbiota.

## Discussion

Our analysis demonstrates that the human gut microbiota’s role in maintaining basal inflammatory homeostasis is a sex-specific process that is profoundly disrupted by oral contraceptives. The total loss of microbial-immune correlations in OC users suggests a hormonal decoupling of the holobiont, where exogenous steroids override the ancestral dialogue between commensal metabolites and host immune sentinels.

We observed instances where aggregate data suggested a positive correlation, yet stratification by biological sex revealed a consistent negative association within each group. This inversion demonstrates that the sexual dimorphism in cytokine baselines and microbial abundance can act as a dominant confounder, creating spurious global trends that contradict the underlying biological interactions. The core methodological contribution of this study is the demonstration that aggregate analyses in microbiome-immunology research can be fundamentally flawed. By uncovering cases of sign reversal (e.g., *Butyricimonas virosa* and *Akkermansia muciniphila*), we show that sexual dimorphism can lead to Simpson’s Paradox, where a true biological signal is masked or cancelled out when sexes are pooled^22,30^. This finding mandates a shift toward sex-stratified models to ensure biological accuracy in future clinical trials and functional studies.

A recurring theme in our results is the functional switch triggered by the presence of serum. Taxa such as *Lawsonibacter asaccharolyticus*, and *Agathobaculum butyriciproducens* transitioned from neutral or slightly pro-inflammatory associations in serum-free media to potent anti-inflammatory roles (inhibiting TNF-*α*) in supplemented conditions. This suggests that the immunomodulatory potential of the microbiota is not intrinsic to the bacteria alone but depends on systemic co-factors present in the host’s circulation. Given that *L. asaccharolyticus* has been previously linked to coffee consumption and anti-inflammatory properties in clinical cohorts^31,32^, our data supports its role as a beneficial commensal in healthy individuals.

The data suggests that males and females may rely on different microbial anchors to maintain systemic quiescence. The finding that *B. ovatus* acts as a negative regulator of basal TNF-*α* levels aligns with its known capacity to induce secretory IgA and promote immune tolerance through complex polysaccharide metabolism^33,34^. Further, male homeostasis appears heavily influenced by butyrate-producers (Roseburia and Alistipes genera) and Lawsonibacter, with a notable role for Akkermansia in priming the myeloid response through IL-6. Meanwhile, on the other side, female homeostasis exhibits a more complex, multi-target regulatory landscape, such as *Firmicutes bacterium CAG:94, Faecalitalea cylindroides* and *Clostridium disporicum*.

Our finding regarding the unannotated Firmicute align with recent evidence^35^, which identified specific ABC-transporters from a Firmicutes with 96% sequence identity to that of CAG:94 capable of exporting muropeptide precursors with potent immunomodulatory effects. The divergent correlation profile we observed –characterized by a reduction in TNF-*α* alongside an increase in IL-22– suggests that *Firmicutes bacterium CAG:94* may function as a ‘homeostatic rheostat’^15^.

By extending our observations to 7 days in PBMCs, we captured the transition from innate ‘sensing’ to adaptive ‘priming’. The consistent association of *Anaerobutyricum hallii* and *L. asaccharolyticus* with Th17 and Th22-related cytokines (IL-17, IL-22) suggests that these commensals are not merely suppressing inflammation^32^, but actively training the adaptive immune system to maintain barrier integrity. A key aspect for interpreting the observed sexual dimorphism is the existence of the gut-vaginal-immune axis. Unlike men, the female immune system is exposed to a second massive source of microbial antigens through the reproductive tract. This dual exposure suggests that female immune cells (such as the macrophages and PBMCs analyzed here) possess a distinctly programmed activation threshold and receptor network. The higher density of microbiome-cytokine interactions observed in NOC women compared to men may reflect the distinct regulatory influence of fluctuating sex hormones on TLR signaling and cytokine production^36^

These results, such as the triple inhibition orchestrated by *Ruminococcus sp. CAG:563* in women, could reflect an evolutionary adaptation to maintain systemic tolerance in the face of the variability of female mucosal reservoirs. While the male system appears to respond more linearly to classic intestinal commensals, the female system shows more nuanced regulation, possibly influenced by the strolome (the community of bacteria that metabolize estrogens) and the need to avoid deleterious inflammatory responses during hormonal cycles. This ‘regulatory redundancy’ in women could explain why certain species, such as *Butyricimonas virosa*, show divergent associations between sexes: the female immune system may be interpreting the signals from this bacterium in the context of a much more complex hormonal and mucosal environment.

By identifying specific microbial species that function as homeostatic anchors^23–25,33,37–44^, the present study provides a functional framework to annotate the roles in human immune regulation of taxa whose immunological signatures remain largely uncharacterized, such as *Butyricimonas virosa, Clostridium disporicum, Faecalitalea cylindroides* and *Ruminococcus sp. CAG:563* among many others. These novel microbiota-cytokine links suggest that the basal inflammatory tone is not a static state, but a finely tuned dialogue susceptible to hormonal and environmental interference. Consequently, these findings open new avenues for mechanistic research and the development of targeted nutraceuticals or microbial-based therapies aimed at restoring immune balance in chronic inflammatory conditions^45–48^.

Despite the robustness of the identified associations, several limitations should be considered. First, our study utilizes a cross-sectional design from the 500FG cohort, which precludes definitive causal inferences regarding the gut-immune axis. Longitudinal studies would be necessary to establish the temporal dynamics of microbial shifts following hormonal intervention. Second, the OC subgroup was analyzed as a single entity; however, different formulations (e.g., varying progestin types or ethinylestradiol dosages) may exert differential effects on immune signaling. Finally, while our residual-based framework successfully identifies homeostatic anchors, these links remain associative and warrant further mechanistic validation through controlled *in vitro* or *in vivo* models.

## Methods

### Human cohort and data integration

This study utilizes publicly available gut microbiota and cytokine data from the Human Functional Genomics Project (HFGP), specifically focusing on the 500 Functional Genomics (500FG) cohort. This cohort consists of healthy individuals of Western European genetic background, all aged 18 years or older. The dataset integrates whole-genome genotypes, fecal microbiome profiles derived from whole-genome shotgun sequencing, and comprehensive cytokine response profiles. Fecal microbiome data were available for 471 individuals, from which relative abundances of 490 gut microbial taxa were extracted. Cytokine data included levels of IL-1*β*, TNF-*α*, IL-6, IFN-*γ*, IL-17, and IL-22, measured across various stimuli. For the purposes of this study, only two basal conditions were considered: RPMI (serum-free) and RPMI supplemented with serum. Stringent exclusion criteria were applied: 45 volunteers were removed due to medication use, non-European descent, or pre-existing conditions (e.g., kidney disease, diabetes mellitus). Additionally, intrinsic host factors including age, gender, body mass index (BMI), and oral contraceptive (OC) use were incorporated into the analysis. After excluding individuals with incomplete data across microbiota, cytokines, and intrinsic factors, a final analytical cohort of 452 healthy subjects was established. This group comprised 195 males and 257 females, the latter including a subgroup of 139 oral contraceptive users. In this study, missing data were managed using pairwise deletion, and no imputation methods were applied.

### Statistical analysis and covariate adjustment

To identify robust associations between gut microbial taxa and cytokine production while controlling for potential confounders, we developed a multi-step statistical pipeline. First, we filtered microbial species based on a prevalence threshold, retaining only those present in more than 3% of the total cohort. For each stimulus-cytokine pair, we constructed an Ordinary Least Squares (OLS) regression model where the cytokine level was the dependent variable, and age, sex, and BMI were included as independent covariates, while OC use was included exclusively as a covariate for the female cohort analysis.

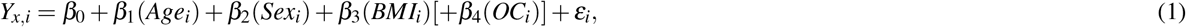

where *Y*_*x,i*_ is the level cytokine *x* of individuo *i. β*_0_ is the intercept and *β*_1_, *β*_2_, *β*_3_, *β*_4_ are the regression coefficients. Age, Sex, BMI, OC: are the covariables. Note that as sex and OC are categoric variables, are leading in the model as dummy variables (0 o 1). *ε*_*i*_ is the residue; it is the most important term for our subsequent analysis. It represents the cytokine variability that could not be explained by age, sex, BMI, or OC. We then calculated the Spearman-rank correlation coefficients *R* between these residuals and the relative abundance of each microbial species. To account for multiple hypothesis testing, *p*-values were adjusted using the Benjamini-Hochberg procedure localized to each stimulus. Only associations with a False Discovery Rate (*q*-value) below 0.05 were considered statistically significant.

### Detection of spurious trends and simpson’s paradox

To ensure the biological authenticity of the identified associations, each significant correlation was subjected to a sensitivity analysis through sex-stratified modeling. We specifically screened for occurrences of Simpson’s paradox, a statistical phenomenon wherein a spurious trend emerges in the aggregate population but is neutralized or inverted upon stratification by confounding variables. In our case, this effect occurs when the overall Spearman-rank correlation coefficient (*R*) exhibits a significant trend that disappears or is opposite to the coefficients obtained within the male and female subgroups. To detect it, besides the regression model (1) we implement a model without sex as a covariate:

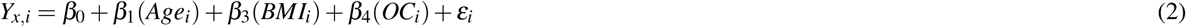

for each gender subgroup, where the cytokine level was the dependent variable, and age, BMI, and OC were included as independent covariates. Then we calculate the respective Spearman-rank coefficients between these residuals and the relative abundance of each microbial species. When such an inversion was detected, the global *R* can be considered an artifact of the differing group means. Furthermore, to decouple the specific effect of OC use, a similar stratification was performed within the female cohort, comparing OC users and non-users while adjusting for age and BMI.

### Network construction and visualization

To integrate and visualize the multi-layered landscape of microbiota-cytokine interactions, we constructed association networks based on the significant correlations identified in the stratified analyses. Nodes in the network represent either microbial species or cytokines, while edges represent significant Spearman-rank correlations (*q <* 0.05). The networks were generated using the Cytoscape Web JS library (v1.0.4)^49^. To distinguish the nature of the associations, edges were color-coded according to the sex or OC use stratification. Dashed and solid edges indicate the RPMI and the RPMI+serum conditions, respectively. To denote the nature of the association without implying a regulatory direction, edges are capped with specific markers: disks represent negative correlations, while arrows represent positive correlations. The layout was optimized using a force-directed algorithm to highlight central homeostatic anchors and the differential connectivity. This topological approach allowed for the identification of cluster-specific signatures on the gut-immune associative density.

## Acknowledgements

I am grateful to the investigators who made their original data available through public repositories. We thank M. Ilid and C. Muglia for critical reading and comments on the manuscript.

## Author contributions statement

L.D. scripting, conception, design of the study and writing of the manuscript.

## Data availability

The datasets analysed during the current study and scripts used are available in the Zenodo repository, DOI:10.5281/zenodo.19120561, https://zenodo.org/records/19120561.

## References

1. Blaser, M. J. & Falkow, S. What are the consequences of the disappearing human microbiota? Nat. Rev. Microbiol. 7, 887–894 (2009).

2. Jiao, Y., Wu, L., Huntington, N. D. & Zhang, X. Crosstalk between gut microbiota and innate immunity and its implication in autoimmune diseases. Front. immunology 11, 282 (2020).

3. Kaoutari, A. E., Armougom, F., Gordon, J. I., Raoult, D. & Henrissat, B. The abundance and variety of carbohydrate-active enzymes in the human gut microbiota. Nat. Rev. Microbiol. 11, 497–504 (2013).

4. Clemmensen, C. et al. Gut-brain cross-talk in metabolic control. Cell 168, 758–774 (2017).

5. Du, Y., Gao, X.-R., Peng, L. & Ge, J.-F. Crosstalk between the microbiota-gut-brain axis and depression. Heliyon 6 (2020).

6. Rhee, S. H., Pothoulakis, C. & Mayer, E. A. Principles and clinical implications of the brain–gut–enteric microbiota axis. Nat. reviews Gastroenterol. & hepatology 6, 306–314 (2009).

7. Wojno, E. D. T. & Artis, D. Innate lymphoid cells: balancing immunity, inflammation, and tissue repair in the intestine. Cell host & microbe 12, 445–457 (2012).

8. Wu, H.-J. & Wu, E. The role of gut microbiota in immune homeostasis and autoimmunity. Gut microbes 3, 4–14 (2012).

9. Mendes, V., Galvão, I. & Vieira, A. T. Mechanisms by which the gut microbiota influences cytokine production and modulates host inflammatory responses. J. Interf. & Cytokine Res. 39, 393–409 (2019).

10. Maynard, C. L., Elson, C. O., Hatton, R. D. & Weaver, C. T. Reciprocal interactions of the intestinal microbiota and immune system. Nature 489, 231–241 (2012).

11. Belkaid, Y. & Hand, T. W. Role of the microbiota in immunity and inflammation. Cell 157, 121–141 (2014).

12. Markowiak, P. & Śliżewska, K. Effects of probiotics, prebiotics, and synbiotics on human health. Nutrients 9, 1021 (2017).

13. Fakruddina, M., Shishirb, M. A., Yousufa, Z. & Khanc, M. S. S. Next-generation probiotics-the future of biotherapeutics. Microb. Bioact. 5, 156–163 (2022).

14. Hou, K. et al. Microbiota in health and diseases. Signal transduction targeted therapy 7, 135 (2022).

15. Orsini Delgado, M. L., Gamelas Magalhaes, J., Morra, R. & Cultrone, A. Muropeptides and muropeptide transporters impact on host immune response. Gut Microbes 16, 2418412 (2024).

16. Cleophas, M. C. et al. Effects of oral butyrate supplementation on inflammatory potential of circulating peripheral blood mononuclear cells in healthy and obese males. Sci. reports 9, 775 (2019).

17. Chen, R. et al. Pattern recognition receptors: function, regulation and therapeutic potential. Signal Transduct. Target. Ther. 10, 216 (2025).

18. Di Vincenzo, F., Del Gaudio, A., Petito, V., Lopetuso, L. R. & Scaldaferri, F. Gut microbiota, intestinal permeability, and systemic inflammation: a narrative review. Intern. emergency medicine 19, 275–293 (2024).

19. Ter Horst, R. et al. Host and environmental factors influencing individual human cytokine responses. Cell 167, 1111–1124 (2016).

20. Li, Y. et al. A functional genomics approach to understand variation in cytokine production in humans. Cell 167, 1099–1110 (2016).

21. Schirmer, M. et al. Linking the human gut microbiome to inflammatory cytokine production capacity. Cell 167, 1125–1136 (2016).

22. Hernán, M. A., Clayton, D. & Keiding, N. The simpson’s paradox unraveled. Int. journal epidemiology 40, 780–785 (2011).

23. Iida, N. et al. Commensal bacteria control cancer response to therapy by modulating the tumor microenvironment. Science 342, 967–970 (2013).

24. Patterson, A. M. et al. Human gut symbiont roseburia hominis promotes and regulates innate immunity. Front. Immunology 8, 1166 (2017).

25. Bae, M. et al. Akkermansia muciniphila phospholipid induces homeostatic immune responses. Nature 608, 168–173 (2022).

26. Bersudsky, M. et al. Non-redundant properties of il-1α and il-1β during acute colon inflammation in mice. Gut 63, 598–609 (2014).

27. Kuhn, K. A., Manieri, N. A.Liu, T.-C. & Stappenbeck, T. S. Il-6 stimulates intestinal epithelial proliferation and repair after injury. PloS one 9, e114195 (2014).

28. Quinn, K. M. et al. Temporal changes in blood oxidative stress biomarkers across the menstrual cycle and with oral contraceptive use in active women. Eur. J. Appl. Physiol. 121, 2607–2620 (2021).

29. Badenhorst, C., Govus, A. & Mündel, T. Does chronic oral contraceptive use detrimentally affect c-reactive protein or iron status for endurance-trained women? Physiol. Reports 11, e15777 (2023).

30. Bonovas, S. & Piovani, D. Simpson’s paradox in clinical research: a cautionary tale. J. Clin. Medicine 12, 1633 (2023).

31. Manghi, P. et al. Coffee consumption is associated with intestinal lawsonibacter asaccharolyticus abundance and prevalence across multiple cohorts. Nat. Microbiol. 9, 3120–3134 (2024).

32. He, Y. et al. Effect of coffee, tea and alcohol intake on circulating inflammatory cytokines: a two sample-mendelian randomization study. Eur. J. Clin. Nutr. 78, 622–629 (2024).

33. Fultz, R. et al. Unraveling the metabolic requirements of the gut commensal bacteroides ovatus. Front. Microbiol. 12, 745469 (2021).

34. Ihekweazu, F. D. et al. Bacteroides ovatus promotes il-22 production and reduces trinitrobenzene sulfonic acid–driven colonic inflammation. The Am. journal pathology 191, 704 (2021).

35. Liuu, S. et al. Identification of a muropeptide precursor transporter from gut microbiota and its role in preventing intestinal inflammation. Proc. Natl. Acad. Sci. 120, e2306863120 (2023).

36. Klein, S. L. & Flanagan, K. L. Sex differences in immune responses. Nat. Rev. Immunol. 16, 626–638 (2016).

37. Liu, L. et al. Bacteroides vulgatus attenuates experimental mice colitis through modulating gut microbiota and immune responses. Front. immunology 13, 1036196 (2022).

38. Leser, T. & Baker, A. Bifidobacterium adolescentis–a beneficial microbe. Benef. microbes 14, 525–551 (2023).

39. Gavzy, S. J. et al. Bifidobacterium mechanisms of immune modulation and tolerance. Gut Microbes 15, 2291164 (2023).

40. Beresford-Jones, B. S. et al. Enterocloster clostridioformis protects against salmonella pathogenesis and modulates epithelial and mucosal immune function. Microbiome 13, 61 (2025).

41. Pujo, J. et al. Bacteria-derived long chain fatty acid exhibits anti-inflammatory properties in colitis. Gut 70, 1088–1097 (2021).

42. Kang, H. et al. Induction of th1 cytokines by leuconostoc mesenteroides subsp. mesenteroides (kctc 3100) under th2-type conditions and the requirement of nf-κb and p38/jnk. Cytokine 46, 283–289 (2009).

43. Qiao, S. et al. Gut parabacteroides merdae protects against cardiovascular damage by enhancing branched-chain amino acid catabolism. Nat. Metab. 4, 1271–1286 (2022).

44. Zhao, C.-N. et al. Phocaeicola vulgatus induces immunotherapy resistance in hepatocellular carcinoma via reducing indoleacetic acid production. Cell Reports Medicine 6 (2025).

45. Basso, P. J., Câmara, N. O. S. & Sales-Campos, H. Microbial-based therapies in the treatment of inflammatory bowel disease–an overview of human studies. Front. pharmacology 9, 1571 (2019).

46. Oka, A. & Sartor, R. B. Microbial-based and microbial-targeted therapies for inflammatory bowel diseases. Dig. Dis. Sci. 65, 757–788 (2020).

47. Kadyan, S. et al. Microbiome-based therapeutics towards healthier aging and longevity. Genome Medicine 17, 75 (2025).

48. Elazzazy, A. M. et al. Where biology meets engineering: Scaling up microbial nutraceuticals to bridge nutrition, therapeutics, and global impact. Microorganisms 13, 566 (2025).

49. Ono, K. et al. Cytoscape web: bringing network biology to the browser. Nucleic Acids Res. gkaf365 (2025).

